# The multiplex network of human diseases

**DOI:** 10.1101/100370

**Authors:** Arda Halu, Manlio De Domenico, Alex Arenas, Amitabh Sharma

**Affiliations:** Channing Division of Network Medicine, Brigham and Women’s Hospital, Harvard Medical School, Boston, MA 02115; Departament d’Enginyeria Informàtica i Matemàtiques, Universitat Rovira i Virgili, 43007 Tarragona, Spain

## Abstract

Untangling the complex interplay between phenotype and genotype is crucial to the effective characterization and subtyping of diseases. Here we build and analyze the multiplex network of 779 human diseases, which consists of a genotype-based layer and a phenotype-based layer. We show that diseases with common genetic constituents tend to share symptoms, and uncover how phenotype information helps boost genotype information. Moreover, we offer a flexible classification of diseases that considers their molecular underpinnings alongside their clinical manifestations. We detect cohesive groups of diseases that have high intra-group similarity at both the molecular and the phenotypic level. Inspecting these disease classes, we demonstrate the underlying pathways that connect diseases mechanistically. We observe monogenic disorders grouped together with complex diseases for which they increase the risk factor. We propose potentially new disease associations that arise as a unique feature of the information flow within and across the two layers.

## Introduction

The advent of next generation sequencing (NGS) and genome-wide association studies (GWAS) has led to the accumulation of a vast amount of disease-gene associations^1^. In addition, high throughput experimental studies and proteomic technologies have resulted in extensive protein interactions maps. Connecting disease related phenotypes to their underlying molecular mechanisms and genetic constituents is crucial for a better understanding of complex human diseases. The emerging field of network medicine offers the tools of network science for distilling relevant insight from the growing sets of molecular disease omics data^2^. One of the earliest attempts at exploring the higher-level implication of disease-gene associations from the network perspective was the construction of genotype–based disease networks, useful to show the global organization of diseases around functional modules^3^ and to infer comorbidity relations between diseases^4^. On the same basis, phenotypic–based disease networks were constructed by text-mining large-scale Medicare data, systematically classifying diseases based on phenotype similarity^5^, facilitating the identification of patterns of disease progression^6^. Since these pioneering works, many studies have focused on adding to the growing compendium of disease-disease associations. For example, Suratanee et al. identified disease-disease associations using a scoring method based on random walk prioritization in the protein-protein interaction network and identified novel disease-disease interactions^7^. Yang et al. measured disease similarity based on differential coexpression analysis to elucidate dysfunctional regulatory mechanisms and arrived at novel interactions between diseases, whose shared molecular mechanisms have recently been uncovered^8^. Menche et al. identified common mechanistic pathways between diseases by the overlap of disease modules^9^.

Despite these large-scale efforts, the characterization of human disease is incomplete if a single source of information, whether molecular or clinical, is considered in isolation owing to the deeply entangled and causal nature of these different types of data. To address this aspect, researchers have started exploring disease associations using multiple data sources. In a recent study, Zitnik et al. applied a matrix factorization based data fusion approach on different molecular and ontological data that resulted in a multi-level hierarchy of disease classification and predicted previously unknown disease-disease associations^10^. In a similar vein, Moni et al. developed a multiplex network model which combines patient specific diagnostics, integrative omics, and clinical data to generate comorbidity profiles of diseases to help stratify patients and potentially derive personalized medicine solutions in the future^11^. Cheng et al. presented a method that simultaneously uses functional and semantic associations to calculate the similarity between pairs of diseases and predict new associations^12^. Sun et al. developed a combined similarity score using annotation-based, function-based, and topology-based disease similarity measures, compared their predictions against genome–wide association studies, and predicted novel disease associations^13^.

The idea of incorporating multiple sources of information finds its direct counterpart in the literature of multiplex and multilayered networks^14^ where it has been shown on various social, technological and biological network^s15-22^ that their structure and dynamics are better understood in terms of multiple interconnected layers of network^s23 24^. Inspired by this prior work, we hypothesize that we should be able to uncover novel information about disease-disease relationships by incorporating genotypic and phenotypic information simultaneously. In this work, we build a multiplex disease network consisting of a genotype–based layer and a phenotype–based layer. We first apply a recently proposed community detection algorithm^25^ on our disease multiplex. Next, we identify cohesive multiplex disease communities to uncover associations between seemingly disparate diseases that have common molecular mechanisms.

The development of a molecular-based disease classification that links genotype and phenotype layers is remarkably challenging and currently remains an unresolved problem. Our method helps resolve this issue by offering an in-depth understanding of cohesive multiplex disease communities. In the age of the proliferation of high-throughput omics methods, disease classification based only on clinical traits and pathological examination is insufficient by itself and may be misleading^26^. The inclusion of multi-omics information in molecular-level disease-disease relations is expected to improve disease classification^26^. Some of the recent disease classification efforts focused on probabilistic clustering algorithms. Hamaneh et al. devised a cosine-similarity approach based on information flow on disease-protein networks, which outputs clusters of similar disease^s27,28^. The grouping of diseases based on their progression (disease trajectory) is another important aspect of disease classification as it presents us with the possibility of predicting future diseases given the patient’s history. Jensen et al. studied the temporal progression of diseases using large-scale health registry data and identified significant classes of trajectories as well as teased out the key diagnoses central to these trajectories^29^. Furthermore, it has been observed that Mendelian disorders often predispose patients to complex disorders. In this respect, an important linkage has been identified regarding the reconciliation of Mendelian and complex disorders, where Blair et al. mined medical record data to infer associations between these two types of disorders and uniquely mapped each complex disorder to a collection of Mendelian disorders^30^. In light of such recent developments, we focus on disease classification as a primary application of our method. We, therefore, deploy our information flow compression community detection technique on the multiplex network of diseases for a proof-of-concept study on disease classification, and analyze our disease communities for biological relevance and novelty.

## Results

### Construction of the multiplex network of human diseases

We used two different data sources concerning genotypic and phenotypic information about diseases. Relationships due to genotype build a gene-disease bipartite network that is projected onto the disease component to obtain the first disease-disease interaction network. Relationships due to phenotype build a symptom-disease bipartite network that is again projected onto the disease component to obtain the second disease-disease interaction network. In Fig. 1 we show a sketch of this procedure applied on a sub-sample of the combined dataset. The multilayer structure is obtained by considering the two projected networks as layers of a multiplex system in which a disease may or may not exist on both layers. The final multiplex network consists of 779 non-isolated diseases with 1,115 genotypic and 5,005 phenotypic relationships (see Methods for further details).

**Figure 1:**
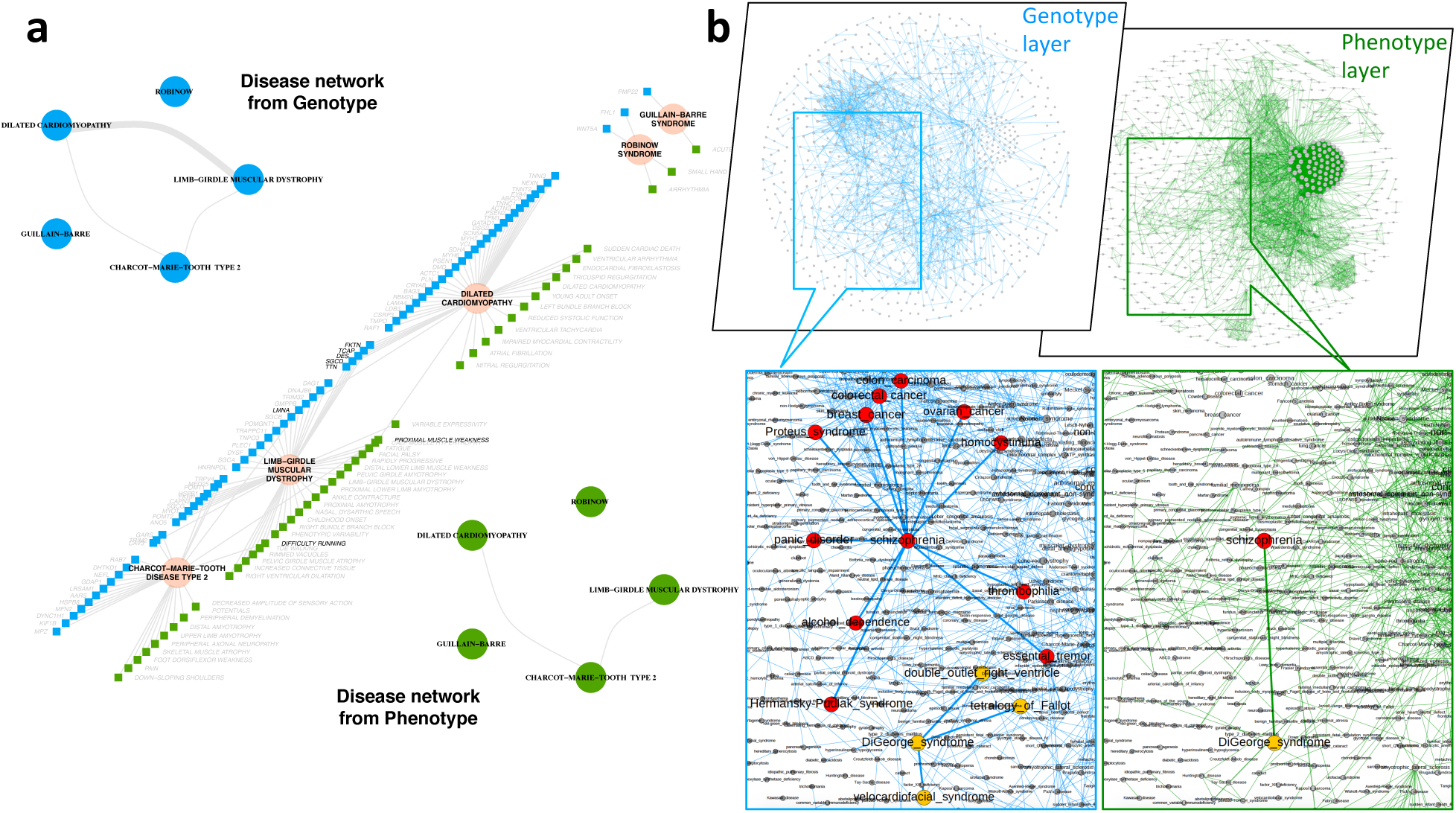
The multiplex disease network. (**a**) Two bipartite networks of disease-gene and disease-symptom interactions are projected onto diseases where diseases are connected in the genotype layer (blue) if they share a common gene and connected in the phenotype layer (green) if they share a symptom. (**b**) Multiplex network detail: Schizophrenia (red) and DiGeorge syndrome (yellow) share genes with 11 and 3 other diseases, respectively, but not with each other. Their connection in the multiplex network is through the phenotype layer, i.e. their shared symptoms.

### Disease-disease interactions shared by genotype and phenotype layers

In order to obtain initial insights about the common genetic and phenotypic mechanisms underlying our disease networks, we considered the shared edges between the genotype and phenotype layer simply when the two network layers are overlaid, i.e. disease pairs that have at least one gene and one symptom in common. Overall, we find that 139 edges overlap between the two layers (Supplementary Figure S1), significantly higher than what would be expected at random (31.80 ± 5.15 edges). The randomization scheme is such that we keep the degree distribution of each network fixed while randomizing the edges per layer for a total number of 5000 times. The distribution resulting from this null model is normal (Shapiro-Wilk test P=0.33) and the z-score for the observed overlap is 20.8 (Supplementary Figure S2). This highly statistically significant enrichment of coinciding disease-disease interactions reveals that diseases that have common genes also tend to share symptoms. Indeed, a closer look at these overlapping edges reveals that they connect disorders that are variable expressions of the same mutation such as MASA syndrome and X-linked hydrocephalus, diseases related to the same gene with important overlapping clinical features such as cerebreal amyloid angiopathy and Alzheimer’s, similar but milder disorders caused by the same gene such as Roberts Syndrome and SC Phocomelia Syndrome, subtypes of a disease such as Gilbert’s Syndrome and Crigler-Najjar Syndrome, as well as subtler associations such as scapuloperoneal myopathy and hypertrophic cardiomyopathy, bronchiectasis and cystic fibrosis, type 2 diabetes mellitus and maturity-onset diabetes of the young, and Noonan syndrome and juvenile myelomonocytic leukemia. It is also interesting to note that on the network-topological level, the different levels of granularity and the distinct local clustering of the two networks (Supplementary Figures S3 and S4) result in the heterogeneous distribution of overlap links around what may be called “overlap hubs” (see Supplementary Figure S5). We find that these hubs are either diseases defined as groups of diseases or multi-system diseases that affect a number of organs and have wide ranging symptoms as well as common genetic factors with other diseases (see Supplementary Information Section 2).

### Gene similarity of disease pairs

An Important indicator of whether or not diseases connected by genotypic or phenotypic relations have a substantially similar genetic background is the gene overlap of disease pairs. To quantify this gene overlap, we calculated the Jaccard index J of the gene sets of diseases connected by an edge. We then calculated the average Jaccard index (J) over all edges in the genotype layer, the phenotype layer, and the edge overlap network, i.e., the network obtained from the multiplex disease network by considering only overlapping disease-disease interactions. We compared the average Jaccard indices of each network with the random expectation, calculated by generating ensembles of networks with the same degree distribution in each layer separately. We observed that the gene similarity of disease pairs in the phenotype network is comparable to random expectation due to the non-specificity of many symptoms, whereas for the genotype layer and the edge overlap network, the gene similarity is significantly different from random expectation (Supplementary Figure S6). Moreover, the gene similarity of the overlap network is higher than both the genotype and phenotype layer. This suggests that the additional layer of information provided by the phenotype layer, despite having little gene overlap overall itself, helps filter out the disease pairs in the genotype layer that have higher gene overlap, reinforcing the genetic effect between disease pairs. In other words, arising as a natural feature of the data, disease pairs whose relationship is supported by both genotype and phenotype evidence have a higher gene overlap than the genotype layer, which hints at the complementary and cooperative nature of these two factors worthy of further investigation.

### Specificity of single-layer disease communities

After building the disease multiplex and probing its structural characteristics, we focus on its higher-level organization. While the simultaneous representation of diseases in the genotype and phenotype space is a very informative abstraction by itself, it is difficult to investigate the interactions between all 779 diseases at once. We, therefore, attempt to decode it further into biologically cohesive disease communities. First, we studied the community partition in each layer separately, i.e., without exploiting the available multiplex information. We identified the communities by using Infomap, a well-known algorithm based on the compression of information flow^31,32^. We selected this method among a number of other community detection methods since it has a direct generalization for multiplex networks (see Supplementary Information Section 5 for a discussion on the choice of community detection technique). As the community detection algorithm does not label communities in any particular way, i.e., it is blind to the underlying biology, we check for the disease overlap between all possible pairs of communities in the genotype and phenotype layers, separately, by calculating the average Jaccard index over all possible pairs. To compare this outcome against a randomized background where the topological aspects of the two networks are conserved, we randomize the network in each layer by keeping the degree distribution constant, using degree-preserving randomization, and appying Infomap on the resulting networks. Once again, we calculate the average Jaccard index for the disease overlap between all pairs of communities, in this case for the randomized ensemble of 5000 realizations. Remarkably, we find that the communities into which the algorithm puts the diseases are layer-specific, meaning that there is little correspondence between the disease communities in the two layers. The average Jaccard index for the disease overlap between layers is 〈J〉 = 0.00286, which is indistinguishable (z-score= 0.882) from that of the degree-preserving randomized layers where the mean of the average Jaccard distribution of the randomized ensemble is 〈J〉 = 0.00261± 0.000289 (Supplementary Figure S7). This indicates that the Infomap algorithm, when applied to genotype and phenotype layers separately, results in distinct disease groupings that are sensitive to the underlying network. However, we argue that disease communities in each layer are meaningful from two different aspects, no matter how distinct they are. In the genotype layer, diseases in the same community represent diseases with common molecular roots, whereas the diseases in the same community in the phenotype layer have similar clinical manifestations. Rather than conflicting pieces of information, we regard these as two complementary sources of information that have to be reconciled using multiplex networks.

### High cohesiveness of multiplex disease communities

We further analyzed the human disease multiplex network to shed light on the functional organization of diseases when genotypic and phenotypic information are considered simultaneously. We calculated multiplex communities by means of Multiplex Infomap^25^, which represents the natural extension of Infomap to multilayer topologies. Given the absence of inter-layer connections between the two layers, the analysis depends on an internal parameter called the relax rate, which regulates the trade-off between random exploration within and across layers (see Supplementary Information Section 5 and Supplementary Figure S8 for a sensitivity analysis for determining the relax rate).

To assess the cohesiveness of the multiplex disease communities that we found using Multiplex Infomap, we look for similarities between disease pairs within communities. Our hypothesis is that if two diseases in a community share clinical characteristics, then they should have common biological pathways and genes. To test this hypothesis, we calculated biological process similarity, comorbidity, gene overlap, and phenotype semantic similarity for each disease pair in a given community. Given that our multiplex consists of a genotype and phenotype layer, we selected these four criteria such that both molecular/genotypic and clinical/phenotypic factors are accounted for. Our goal is to show that multiplex communities are able to capture known disease-disease relationships successfully, as well as offer new insights into unknown disease relationships.

Our corpus of 779 OMIM diseases, consisting of complex as well as monogenic disorders, is divided into 128 multiplex communities, the largest of which has 91 diseases and the smallest of which has 2 diseases. The number of unique diseases classified into groups saturates quickly as we proceed cumulatively from the largest group to the smallest group (Supplementary Figure S9). Here, we concentrate on the 29 largest diseases classes of size 10 and greater for statistical evaluation, which overall comprise 520 diseases.

It is worth noting two unique properties of our multiplex community detection method: namely, a disease can be assigned i) to the same community twice, i.e., it belongs to the same community across the two layers, and ii) to two different communities. In the first case the community structure can be distinguished between the two layers such that one can state that a disease belongs to the same class in both the genotype and the phenotype layer. The second case is especially important for our purposes because it defies the current clinical observation–based disease classification and allows for a more organic classification with more flexible boundaries that respect the molecular underpinnings of diseases. In our dataset, 215 diseases appear in two classes, acting as bridges between those two classes (Supplementary Figures S10 and S11). We note that these “bridge diseases” arise from the overlapping multiplex communities. As a result, the 128 communities have 994 diseases overall instead of 779.

As a general survey of all the communities, we first map the pairwise similarities of the diseases in all communities in a heatmap where we order the communities in decreasing size order (see Supplementary Figures S12-S15). For each community, we assess the significance of the intra-community similarity according to the four metrics previously introduced by comparing the distributions of the community values against the background of all disease pairs in all communities (Figure 2a). We use the Mann-Whitney U test to calculate P-values and consider as significantly similar the communities that have an average similarity higher than the background and a P-value smaller than 0.01. For the 29 largest communities with size greater than or equal to 10, we find that the majority of the communities have significantly high similarity. In particular, the number of communities that have significantly high similarity is 26/29 for comorbidity measured in terms of relative risk (RR), 23/29 for gene overlap measured by Jaccard index, 20/29 for GO: Biological Process similarity, and 27/29 for semantic similarity measured by MimMiner. Furthermore, 14/29 clusters have significantly high similarity with respect to all of the four similarity measures, and 29/29 clusters have significantly high similarity by at least two of the four similarity measures (Figure 2b). Taken together, this shows that owing to the fusion of information on both layers of the disease multiplex, our multiplex disease communities are homogeneous at both molecular and phenotypic levels.

**Figure 2:**
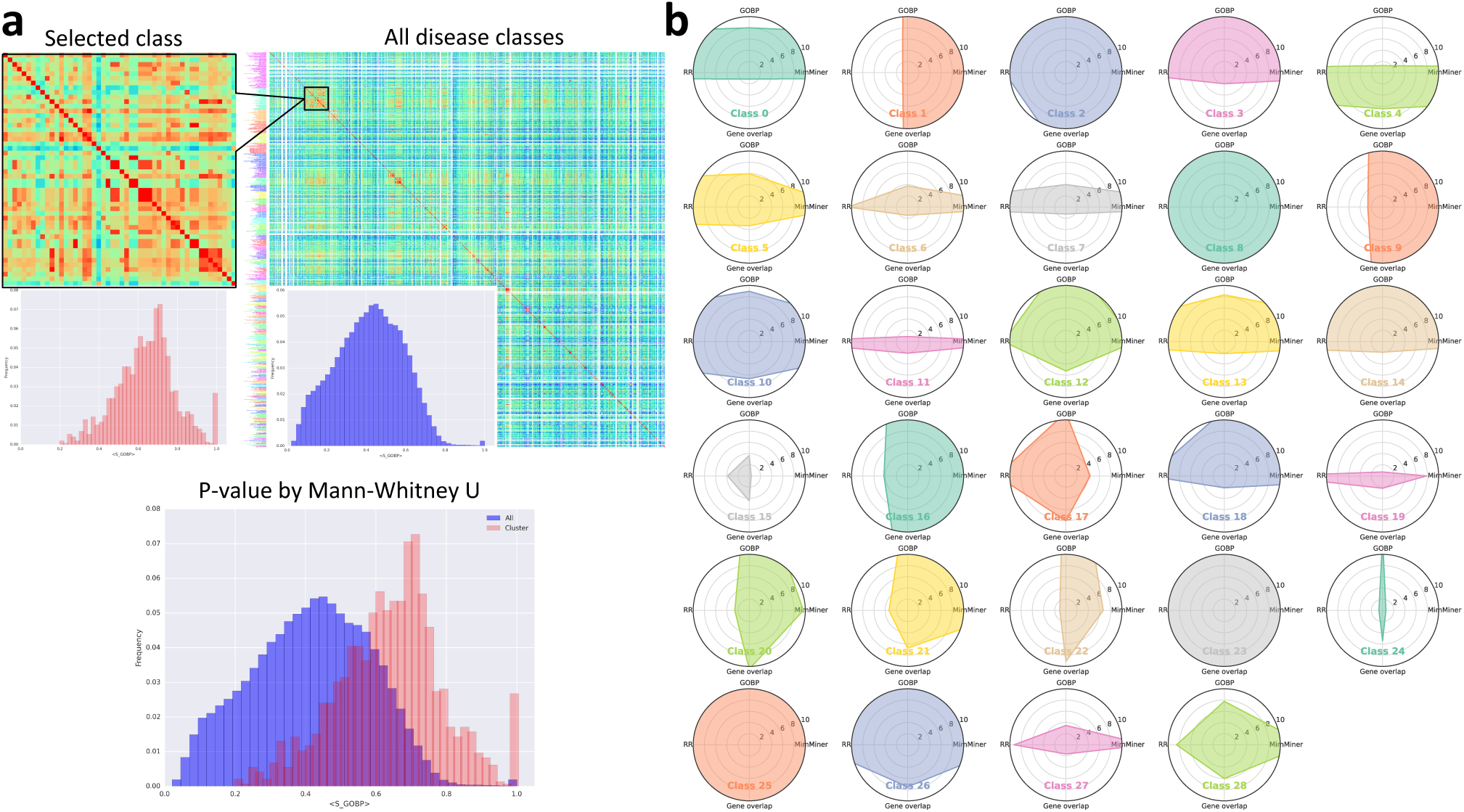
Intra-class similarity assessment. (**a**) Each disease class is evaluated for significance of intra-class similarity according to the four similarity measures. For each similarity measure, the distribution of similarity values for the disease class is compared against the background of all disease classes and its respective P-value is calculated by Mann-Whitney U test. (**b**) Radar plots of the 29 disease classes with size 10 or more, showing the -log P-values. The molecular similarty represented by the North-South axis is for GO:BP and Gene Overlap, whereas phenotypic similarity represented by the East-West axis is for relative risk (RR) comorbidity and MimMiner phenotype semantic similarity. The larger the overall shaded area, the more significant the intra-class similarity. Points inside the innermost circle (-log P-value < 2, or P-value > 0.01) represent non-significant intra-class similarities for the respective similarity measure.

For a further validation of the multiplex disease communities, we verify that they are distinct from randomized communities in that they have significantly higher similarity than random according to the four similarity measures (Supplementary Information Section 9). More importantly, to test the non-additive cooperative effect of multiplexity and to give a meaningful background against which the cohesiveness of multiplex disease communities can be compared, we carried out the same molecular and phenotypic similarity assessment on genotpye and phenotype-layer specific disease communities (Supplementary Information Section 11). We found that, i) multiplex communities perform better than the genotype layer classes in terms of phenotypic similarity measures, and ii) multiplex communities perform better than the phenotype layer classes in terms of the molecular similarity measures while performing comparably with or better than the phenotype layer classes in terms of the phenotype similarity measures. This suggests that the multiplex disease network is indeed “greater than the sum of its parts,” compensating for the lacking features of each single layer and reflecting an all-around cohesive picture rather than only for one of the two aspects.

### Confirming established disease associations and finding new ones

We go one step further into the disease-level and investigate disease relationships in multiplex communities for new biological insight. Our aim is two-fold: first, verify known disease relationships, showing that the communities are reliable, and second, uncover novel disease-disease relationships. To this end, we first select disease pairs that are expected from known literature associations. We calculate the cooccurrence of diseases in the entire PubMed database using the PubAtlas query tool (see Methods) and build a “publication co-occurrence network” of diseases in the multiplex community. This helps us to identify the disease pairs with high co-occurrence, which we would expect to see in the same disease community based on prior research, as opposed to possibly novel disease pairs with few literature co-occurences.

For example, we see that Community 23, which is one of the communities with the highest intra-class similarity, mainly consists of rare, inherited skeletal abnormalities (Figure 3). Among these, some connections are well established in literature (Figure 3e), such as that between the Hunter-Thompson and Grebe types of acromesomelic dysplasia, both of which are caused by a mutation in the GDF5 gene. Similarly, tarsal-carpal coalition syndrome and proximal symphalangism, which are both caused by a heterozygous mutation in the NOG gene, have high co-occurrence in the literature. The close relationship of these expected pairs of diseases are further evidenced by their high comorbidity, high biological process similarity, and high phenotype semantic similarity. Multiple synostoses syndrome also has high publication co-occurrence with tarsal-carpal coalition syndrome and proximal symphalangism, with which it shares one of its three related genes. It has high biological process similarity and moderate phenotype semantic similarity with those two diseases. In addition to these already established relationships, our multiplex disease communities also bring together many diseases with subtler connections. For instance, fibular hypoplasia and complex brachydactyly are also closely associated with acromesomelic dysplasia types, and arise from a mutation in the same gene, GDF5, even though these diseases do not appear together in the literature and have low comorbidity. Furthermore, they have high biological process similarity and moderate semantic similarity with acromesomelic dysplasia types. Likewise, even though synpolydactyly and acrocapitofemoral dysplasia do not have any shared genes, they are grouped together owing to their high comorbidity and similar GO biological process terms. Given that acro-capitofemoral dysplasia is a recently delineated skeletal dysplasia characterized by brachydactyly, synpolydactyly, which is another digit dysplasia, might have common underlying molecular mechanisms with this rare disease.

**Figure 3:**
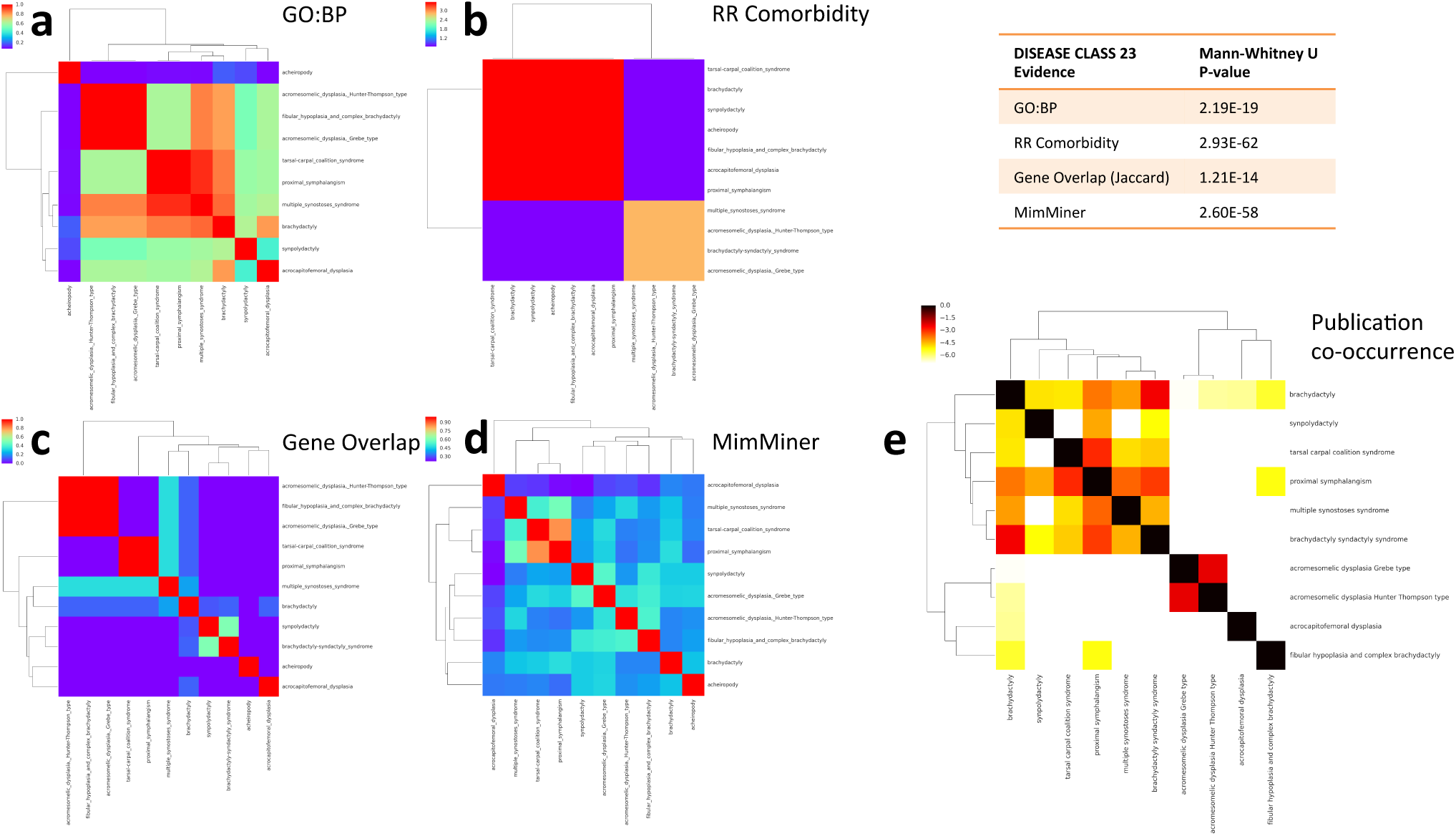
A closer look at the disease classes. Disease Class 23 is characterized by rare skeletal abnormalities. (**a**) GO: Biological Process similarity heatmap, where similarity scores range between 0 and 1. (**b**) Relative risk (RR) comorbidity heatmap, where the colors represent the logarithm of the RR values. (**c**) Gene overlap, quantified by the Jaccard index. (**d**) MimMiner phenotype semantic similarity heatmap, where similarity scores range between 0 and 1. (**e**) PubMed literature co-occurrence heatmap and representative network, where the number in each cell denotes the number of publications that have the co-occurence of the queried keywords. The color of each cell represents the Jaccard index and a redder cell means higher literature co-occurrence weighted by the total number of publications.

These new connections show that our multiplex disease classes are able to capture the molecular basis of connections between diseases even if they have not been associated with each other simply by their clinical manifestations.

By inspecting Community 8, another one of the communities with the highest intra-class similarity, we see that it groups together rare skeletal abnormalities with rare congenital heart defects, which usually have related clinical manifestations as well as shared genetic causes (Figure 4). DiGeorge syndrome and velocardiofacial syndrome have traditionally been studied together in literature, and both are caused by a hemizygous deletion of chromosome 22q11.2 and are also believed to be caused by point mutations on the TBX1 gene. Tetralogy of Fallot, by contrast, is clinically associated with four types of atrial and ventricular defects: atrial septal defect, ventricular septal defect, double outlet right ventricle, and atrioventricular canal defect, classified with the same ICD9 code (745). These diseases also share many genetic elements resulting in them being clustered together in terms of gene overlap (Figure 4c). Overall, we see that DiGeorge syndrome/velocardiofacial syndrome and the cardiac defects related to Tetralogy of Fallot all have very high GO Biological Process similarity despite not sharing many genetic factors (Figure 4a). When we look closer at two of the diseases in this group, namely DiGeorge syndrome and ventricular septal defect, at the molecular level, we see that biological processes related to cardiac development such as outflow tract morphogenesis (GO: 0003151), heart morphogenesis (GO: 0003007), heart development(GO: 0007507), as well as processes related to endocrine development, such as thyroid gland development (GO: 0030878), are shared between the genes of these two diseases, underpinning the biological similarity of the shared symptoms of these diseases. From a developmental biology perspective, these disorders in part reflect dysmorphogenesis at the branchial cleft level, which suggests that developmental defects are driven by otherwise “Mendelian” mutations in a unique (temporal) developmental biology context. To assess the significance of the biological similarity between these disease genes and the related biological process, we apply the GS2 algorithm on a randomly selected genes from the interactome and calculate the biological process similarity with each of the above processes. As expected, both diseases have significantly higher biological process similarity with outflow tract morphogenesis (*S_GOBP_* = 0.687, z-score= 2.57 and *S_GOBP_* = 0.747, z-score= 2.87 for ventricular septal defect and DiGeorge syndrome, respectively), heart morphogenesis (*S_GOBP_* = 0.686, z-score= 2.45 and *S_GOBP_* = 0.746, z-score= 2.72 for ventricular septal defect and DiGeorge syndrome, respectively), heart development (*S_GOBP_* = 0.676, z-score= 2.17 and *S_GOBP_* = 0.727, z-score= 2.41 for ventricular septal defect and DiGeorge syndrome, respectively) and thyroid gland development (*S_GOBP_* = 0.656, z-score= 2.23 and *S_GOBP_* = 0.717, z-score= 2.65 for ventricular septal defect and DiGeorge syndrome, respectively), compared to random expectation (Figure 6). The other important disease group in this community consists of sclerosteosis, craniometaphyseal dysplasia, oculodentodigital dysplasia, syndactyly, chondrocalcinosis 3MC syndrome, and Hajdu-Cheney syndrome, which are all anomalies of the bones. Hajdu-Cheney syndrome, in particular, has many cardiovascular manifestations, including atrial and ventricular septal defects, and is, hence, effectively the common disease that bridges the cardiovascular and bone related diseases in this disease class.

**Figure 4:**
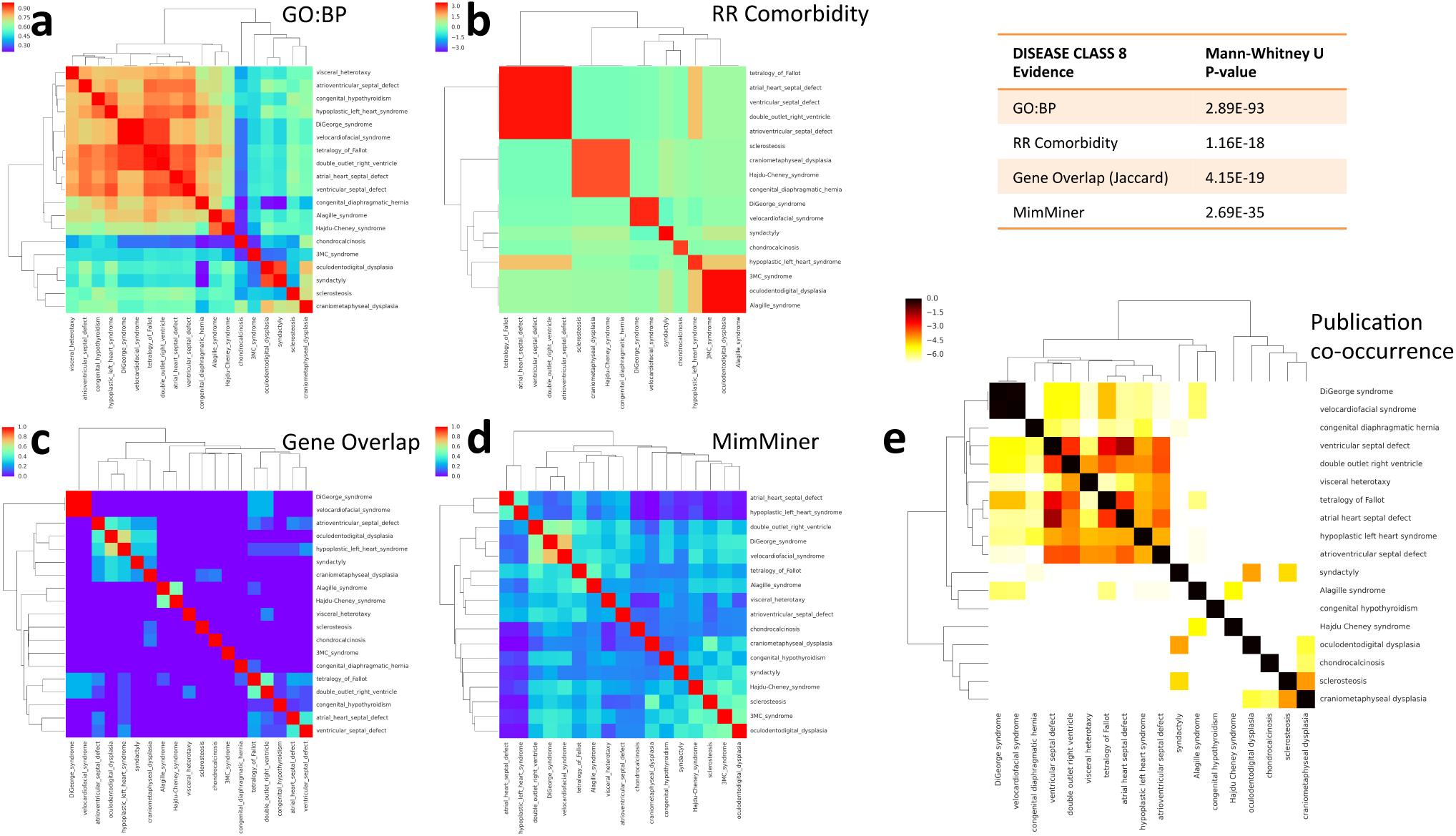
A closer look at the disease classes. Disease Class 8 is characterized by rare congenital heart defects and skeletal anomalies. (**a**) GO: Biological Process similarity heatmap, where simi-larity scores range between 0 and 1. (**b**) Relative risk (RR) comorbidity heatmap, where the colors represent the logarithm of the RR values. (**c**) Gene overlap, quantified by the Jaccard index. (**d**) MimMiner phenotype semantic similarity heatmap, where similarity scores range between 0 and 1. (**e**) PubMed literature co-occurrence heatmap and representative network, where the number in each cell denotes the number of publications that have the co-occurence of the queried keywords. The color of each cell represents the Jaccard index and a redder cell means higher literature co-occurrence weighted by the total number of publications.

**Figure 6:**
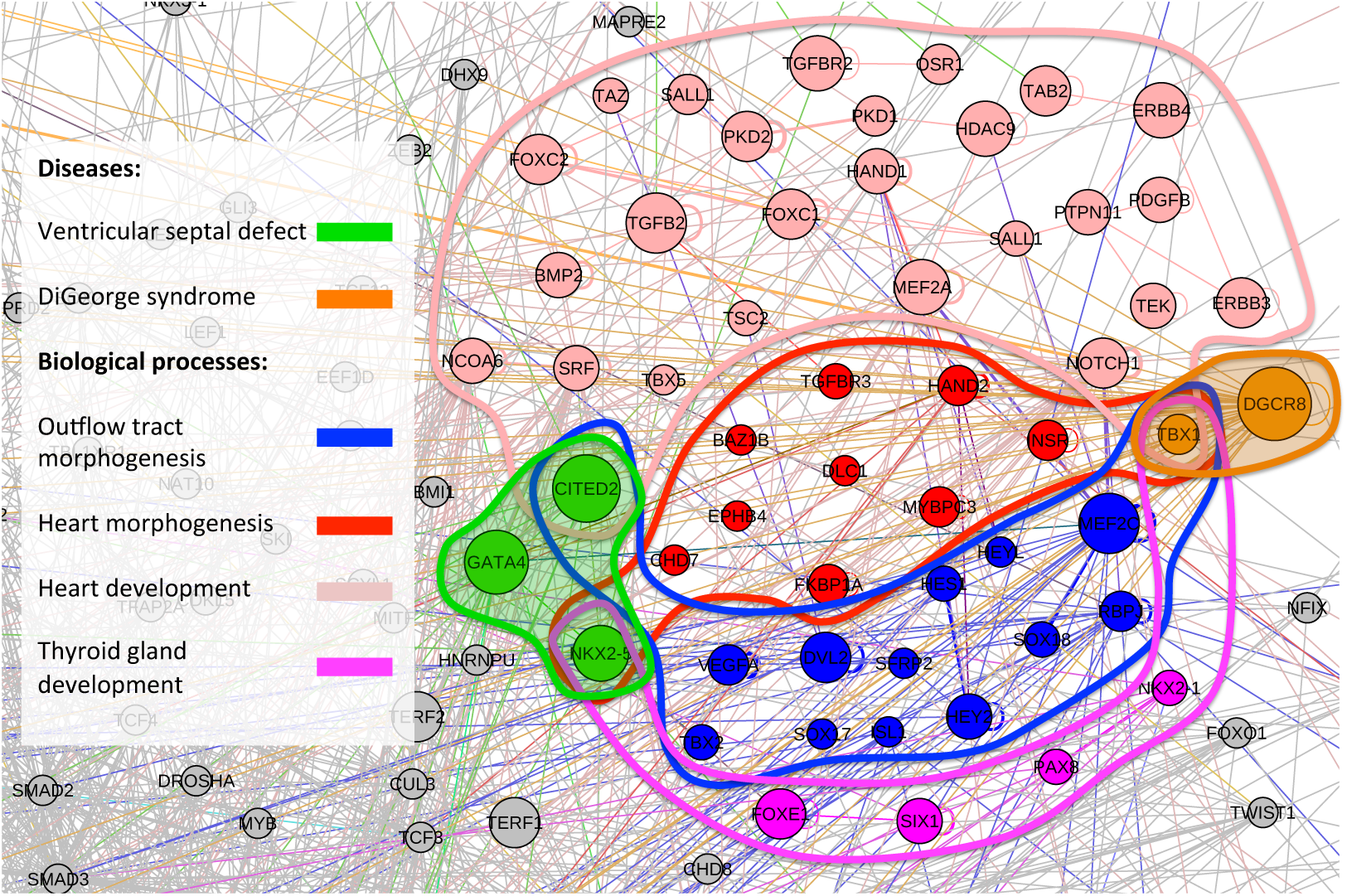
Biological process similarity of diseases. Gene associated with ventricular septal defect (green) and DiGeorge Syndrome (orange) are connected by biological processes related to cardiac development (outflow tract morphogenesis in blue; heart morphogenesis in red; heart development in pink) and endocrine development (thyroid gland development in magenta).

As an additional check to verify the reliability of the boundaries of the multiplex disease classes, we compare the intra-class publication co-occurrences with inter-class publication occurrences (Supplementary Information Section 10). For the two disease classes discussed here, we find that the publication co-occurrence of diseases within Class 8 and Class 23 are significantly higher than the publication co-occurrence of diseases between Class 8 and Class 23 (Supplementary Figure S16). Generalizing this analysis to all multiplex disease classes, we show that while a few classes are relatively more convoluted (meaning, despite having higher intra-class publication co-occurrence than inter-class, have a few non-significant instances) (Supplementary Figure S19), overwhelmingly, the intra-class publication co-occurrence was higher than the inter-class case for all classes (Supplementary Figure S18).

Smaller communities with five or fewer diseases contain a smaller fraction of the overall diseases, with 72 classes comprising 146 diseases. A quick look at some of the many smaller classes reveals that these classes are mostly completely homogeneous, consisting of synonymous diseases, Mendelian diseases and their direct complications, subtypes of the same disease, or diseases with close genetic roots (see Supplementary Table 1). They nevertheless offer very interesting disease associations that are novel. For instance, we see that idiopathic pulmonary fibrosis (IPF), is grouped together with common variable immunodeficiency and immunoglobulin alpha (IgA) deficiency. Indeed, IgA in serum has recently been proposed as a prognostic biomarker for

IPF ^33^. While space limitations preclude us from discussing every disease class in detail, given the high cohesiveness of multiplex disease classes, readers can draw similar possibly interesting disease-disease interactions from any of the classes using Supplementary Table 1 as reference.

Finally, for a complete picture of disease-genotype relations, we have carried out the same analysis on the disease multiplex constructed using Genome Wide Association Studies (GWAS) data for the genotype layer (see Supplementary Information Section 12). Our results show that, once again, despite the distinct sizes and topologies of the GWAS and OMIM disease-gene networks ^34^, we are able to capture cohesive disease groups. As an example, Class 1 (Figure 5) remarkably brings together various types of amyotrophic lateral sclerosis (ALS) with a group of cancer subtypes. The association between ALS and cancer has recently been suggested, where significantly elevated risk of ALS death among survivors of melanoma has been shown^35^.

**Figure 5:**
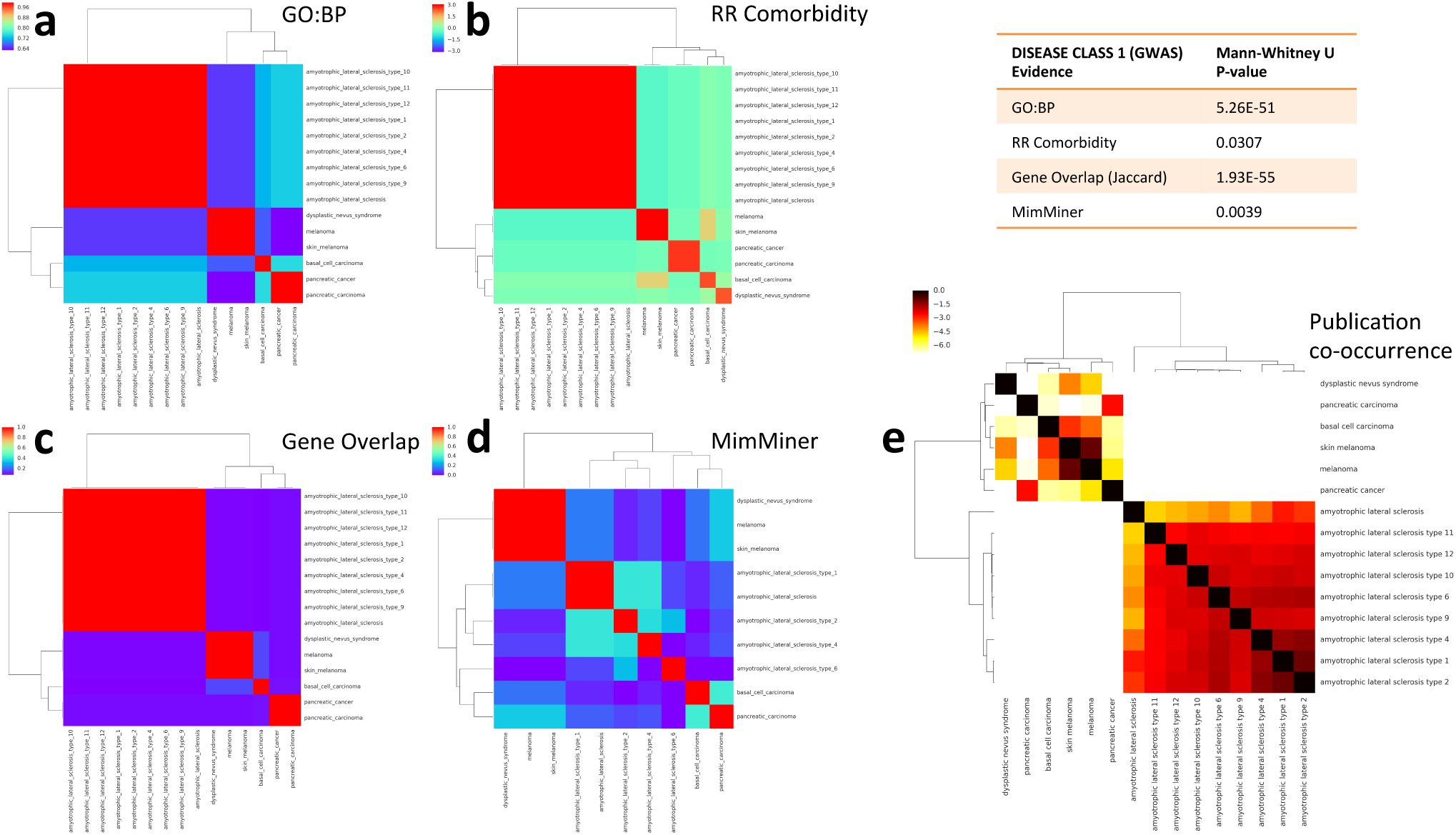
A closer look at the disease classes. Disease Class 1 in the GWAS dataset is character-ized by ALS types and cancers. (**a**) GO: Biological Process similarity heatmap, where similarity scores range between 0 and 1. (**b**) Relative risk (RR) comorbidity heatmap, where the colors rep-resent the logarithm of the RR values. (**c**) Gene overlap, quantified by the Jaccard index. (**d**) MimMiner phenotype semantic similarity heatmap, where similarity scores range between 0 and 1. (**e**) PubMed literature co-occurrence heatmap and representative network, where the number in each cell denotes the number of publications that have the co-occurence of the queried keywords. The color of each cell represents the Jaccard index and a redder cell means higher literature co-occurrence weighted by the total number of publications.

These proof-of-concept examples show that the multiplex network of human diseases, coupled with community detection methods designed to highlight the interplay between different layers of the network, captures the biological similarity implicit in different data sources. It, therefore, brings a new dimension to disease relationships that cannot be achieved if we study disease networks from the genotype and phenotype aspects separately.

## Discussion

To obtain an in-depth understanding of the molecular basis of human disease, we need to recognize the complex relationship between genotype and phenotype in response to environmental and genetic influences. Today, one of the most important factors hampering the effective characterization of diseases is the lack of apparent connection between how they manifest and what their molecular underpinnings are. With this issue as the starting point of this study, we made an attempt to analyze systematically the structural and functional aspects of the multiplex network of human diseases. We showed that diseases with common genes tend to share symptoms, and we uncovered the complementary nature of genotype and phenotype by examining the interplay between the two network layers. We reported our finding that, through the representation of disease relationships in a multiplex network, a wide range of monogenic and complex diseases can be grouped into classes that are cohesive at both molecular and symptomatic levels. We argued that the disease classes found in this way offer a flexible description of pathobiological processes that melds together genotypic and phenotypic aspects, while community detection applied on separate layers results in layer-specific and distinct disease groupings. Here, we find it important to note the inherent effect of the current incompleteness of the genotype and phenotype layers, which stem from many yet-unknown disease-gene or disease-symptom relationships. While we demonstrated the advantage of the multiplex disease network in its present form over single-layer disease networks combined and discovered interesting disease associations, we believe the accuracy of our method will increase in time as more and more of the missing parts of each network are uncovered.

In contrast with the bottom-up “disease module” approach, which starts with the known genetic determinants of diseases to build disease modules and infer disease-disease interactions based on the localization of these modules in the underlying protein-protein interaction (PPI) network^3 9^, we used a top-down approach, where the starting point was diseases themselves rather than the PPI, and recovered the underlying molecular mechanisms that associate those diseases. One advantage of the top-down approach in characterizing diseases is that the construction of the disease-disease network relies on the unambiguous and relatively relaxed criterion of one shared gene or one shared symptom, making in easier to merge and parse large disease-related knowledge bases such as OMIM and Disease Ontology. Overall, it is worthwhile to compare and contrast the top-down and bottom-up approaches in future studies as they are likely to offer complementary benefits and possibly a significant overlap of results. We also recognize the possible limitations of our current “unweighted” network approach, where a link is established between diseases if the minimal criterion of one shared symptom or gene is met regardless of how many symptoms or genes are shared between them, such as the possible over-inclusion of edges due to common symptoms. However, our aim at this initial stage is to work on the unweighted network to serve as a minimal model of disease-disease interactions in the genotype and phenotype layer. While within the scope of this work we are able to validate the resulting multiplex disease groupings through intra-class similarity assessment and the recovery of known disease-disease relationships, we find it important to note that the results can be improved by treating each layer as weighted in future studies.

Our selection of diseases is geared towards maximizing the size of the dataset and is, therefore, biased towards neither complex nor Mendelian diseases. It, rather, includes diseases of a wide range of prevalences and penetrances, from rare congenital diseases to common cancer types and metabolic disorders. Our method, however, seamlessly integrates the two types of diseases, with OMIM and GWAS evidence alike.

At the core of our disease classification is a technique that takes into account the information flow within and across different network layers. In the context of pathobiology, we use the term “information flow” to refer specifically to the process by which distant diseases can “communicate” by propagating genetic and symptomatic signals through intermediary diseases in the global disease network whereby, for instance, disease A, which shares certain genes (or symptoms) with disease B, can be related indirectly to disease C, which also shares certain genes (or symptoms) with disease B. The diseases sending and receiving more flows from each other in the genotype layer would likely have related molecular roots whereas in the phenotype layer, this would mean groups of diseases that share clinical manifestations. We note that the use of quotes in “communication” is in order to emphasize that this is a process that does not involve actual molecular signaling but rather a conceptual one that implies at associations between diseases. Capable of traversing both layers of disease networks, this method can put the same disease into two different classes based on both genotypic and phenotypic data. This “redundancy” of disease classes, with “bridge diseases” connecting them, is a key component of the next generation of organic, flexible disease classifications, where we can develop a network of disease classes rather than the current tree-like, hierarchical classification where a disease can only belong to one class. We believe that the common biology of these bridge diseases deserves further detailed investigation, and note it as an important future direction.

Overall, this study provides a rich compendium of disease associations and groupings that can be examined from many aspects ranging from relationships between complex and Mendelian disorders, to possible commonalities between classes of diseases. To our knowledge, this is the first time relationships between diseases have been assessed from the perspective of multiplex networks using novel network techniques designed specifically to uncover systemic properties at multiple levels.

A notable finding supported by clinical and genetic observations is that rare, Mendelian disorders predispose individuals to more common, complex diseases ^30^. In fact, in many of the disease classes, we observe monogenic disorders grouped together with the complex diseases for which they increase the risk factor. For instance, patients with Denys-Drash Syndrome, which is a rare disorder characterized by abnormal kidney function believed to be due to a mutation in the WT1 gene, have an estimated 90 percent chance of developing a rare form of kidney cancer known as Wilms tumor. Affected individuals may develop multiple tumors in one or both kidneys, testes, or ovaries. We note that this disease, along with Frasier syndrome, is grouped together with cancer types in disease class 16. Similarly, it has been documented that DiGeorge syndrome shows an increased risk of schizophrenia^36^, which are both grouped in disease class 21. Disorders related to the homozygous or compound heterozygous deletions and loss-of-function mutations in NPHP1, such as Joubert Syndrome or Senior-Loken syndrome, have been linked to autistic spectrum disorder^37^, all of which can be found in disease class 12. Individuals affected by Wolfram syndrome have diabetes mellitus and degeneration of the optic nerve, and we note these diseases in class 22. Many more examples in line with the complex-Mendelian disorder associations can be given by inspecting our disease classes. Our disease classes are, therefore, inclusive units wherein complex and Mendelian disorders are linked with both genetic and clinical factors.

Another interesting observation is that some Mendelian diseases with severe phenotypes, which were believed to be completely penetrant but were recently identified to be present in individuals with no apparent clinical manifestations ^38^, were found in our dataset within the smaller disease classes with size < 10 (Cystic fibrosis: class 73(4), Smith-Lemli-Opitz syndrome: Class 40 (7), Pfeiffer syndrome: class 82 (3), atelosteogenesis: class 31(9)). This may point to the fact that these severe childhood diseases, which may potentially have individuals resilient to them, tend to avoid being classified alongside other diseases, but, rather, have their own small cohesion groups. Although the findings of that particular study currently remain to be fully verified, it nevertheless represents an interesting research frontier wherein genetic modifiers that impact phenotypic variability are found through phenotype-genotype correlations across multiplex networks ultimately to identify “fully penetrant” Mendelian diseases that can be harbored by resilient individuals more clearly.

The results of our work can be refined and built upon in many ways. With the increased momentum towards precision medicine, efficient subtyping of diseases becomes a crucial need. To fill this need, multiplex disease networks that include a number of disease subtypes can be analyzed and re-grouped using network techniques. Similarly, a better classification of spectrum disorders, which are diseases that are a collection of multiple diseases, represents an important frontier of disease characterization. From that perspective, complex diseases that are regarded as spectrum disorders can benefit from the new associations and disease groupings that our method provides. Finally, since the field of multiplex networks and the subsequent methods to detect communities within them is a very rapidly evolving one, more sophisticated multiplex community detection methods can further improve the characterization of diseases and identification of disease groups in the future.

## Methods

### The genotype data set

The disease-gene bipartite network is built from the well known Online Mendelian Inheritance in Man (OMIM) data set. OMIM^139,40^ is a knowledge base whose content is derived exclusively from the published biomedical literature describing human phenotypes and genes. The version used in this study contains information about 3,369 genes and 4,239 diseases, for a total of 5,308 edges.

### The phenotype data set

The disease-symptom bipartite network is built from the well known Human Phenotype Ontology data set. Human Phenotype Ontology^41^ provides a structured, comprehensive and well-defined set of 10,088 classes (terms) describing human phenotypic abnormalities and 13,326 subclass relations between the HPO classes. We have found 6,662 diseases with relationships to 11,052 symptoms, for a total of 100,281 edges.

### Linking genotype and phenotype data sets

Among the 3,919 diseases common to both databases, we focused our attention on the subset of diseases matching the Disease Ontology database. Disease Ontology^42^ represents a comprehensive knowledge base of developmental and acquired human diseases. It semantically integrates disease and medical vocabularies through extensive cross mapping and integration of several disease-specific terms and different identifiers. We found 2,255 matching unique OMIM identifiers for 910 diseases. Finally, we discarded the isolated diseases without any links in either layer, which resulted in 779 diseases in the multiplex network.

### Community detection on single and multilayer networks

Infomap is an algorithm to optimize the map equation^31,32^. This information-theoretic equation makes extensive use of the duality between the problem of compressing a data set and the problem of revealing significant structures within it. To achieve its goal, the algorithm makes use of random walkers to explore the network, encoding their flow with sequences of bits. In our case, the data set is a network and the (unknown) significant structures are the communities. Each possible community partition of the network result in a specific encoded flow: the partition whose description length is minimum is the optimal one. Therefore, the Infomap algorithm exploits the dynamics on the network to reveal the underlying community structure.

The generalization of this algorithm to the case of multilayer networks, called Multiplex Infomap^25^, is based on the same principle. In this case, the random walkers explore all the layers of the multilayer structure, and there is a parameter, called the relax rate, regulating the amount of the probability to visit more nodes within the same layer than across layers. In other words, this parameter is used to modulate the “flexibility” of movement across layers in the absence of information on the actual interlayer link weights. Numerical experiments^25^ show that values of the relax rate close to approximately 0.5 are generally good enough to balance exploration within and across layers. We accordingly use a relax rate of 0.45 throughout the analysis.

### Similarity measures: MimMiner

In order to determine the phenotypic similarity of the diseases in a given cluster, we used MimMiner similarity matrix^5^. MimMiner assesses the semantic similarity of phenotypic terms related to diseases in the OMIM database. For each OMIM disease, it builds a feature vector consisting of Medical Subject Headings (MeSH) concepts that collect all synonyms and uniquely identify terms, which makes it a more versatile method than keyword-based searches. The MimMiner similarity score is calculated using these feature vectors, resulting in normalized values between 0 and 1. In our analyses, we used the MimMiner similarity matrix to determine the intra-class similarity of all of the 30 classes of diseases. Of the 779 OMIM diseases we consider in our dataset, 675 (87%) were mapped to the MimMiner matrix. We follow the cutoff of 0.3 proposed in the original paper^5^ to define associations that are biologically informative, whereas we deem scores above 0.6 to be significantly functionally similar.

### Similarity measures: Gene Ontology based on gene set similarity

For an insight into the similarity of the molecular mechanisms underlying the diseases in our disease classes, we make use of the GO-based gene set similarity measure proposed in^43^. We prefer the gene set based similarity over pairwise gene similarity measures since pairwise gene similarity does not scale as well to large gene sets and since we aim to compare pairs of diseases in each class, many of which have multiple genetic elements. This measure essentially lets us rank each GO term with respect to a gene set based on the number of genes in the set that are annotated by the ancestors of that GO term. We limit our attention to Biological Process. Furthermore, for concreteness, we only consider evidences EXP, IPI, IDA, IMP, IGI, IEP, ISS, ISA, ISM and ISO, which are either experimental or computational analysis evidence codes.

### Similarity measures: Comorbidity (Relative Risk)

Another similarity measure we use in assessing the homogeneity of our disease classes is comorbidity, i.e. the co-occurrence of diseases in the same patient. For this, we use a healthcare dataset comprising the patient history of 13 million elderly Americans over the age of 65 covered by the Medicare program^6^. We manually curate our disease set consisting of 779 diseases to reflect the 3-digit ICD-9 disease classification code. We quantify the comorbidity of disease pairs using relative risk (RR) score, which is given by the ratio of the observed co-occurrence of a disease to the expected probability of co-occurrence if the diseases were independent from each other,

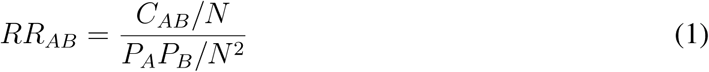
where *C_AB_* is the observed co-occurrence of disease A and B, *P_A_* and *P_B_* are the prevalence of disease A and B, and N is the total number of patients in the dataset’s population. A relative risk greater than 1 indicates a comorbidity that is higher than expected. Since RR values can vary several orders of magnitude depending on the prevalence of a disease, we take the logarithm of RR values when visually representing them.

### Similarity measures: Gene Overlap

As an additional proxy of the molecular intra-similarity of disease classes, we calculate the direct gene overlap between pairs of diseases. This is intended to provide a direct and straightforward measure of shared genetic constituents. We then use Jaccard index J to quantify the gene overlap with

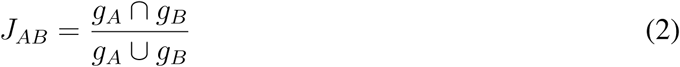
where *g_A_* and *g_B_* are the gene sets of disease A and disease B.

### Publication co-occurrence

To validate the biological meaningfulness of the content of our disease classes, we seek to gain an overview of the literature associations of diseases from PubMed records. PubAtlas^2^ is a web service that acts as a front end to the PubMed/MEDLINE database. Using PubAtlas, we generate “literature heatmap” for each disease class by querying the names of diseases within each class. The color in these heatmaps is given by the logarithm of the Jaccard association value. We also represent this heatmap with a network of literature associations where the color (same as the heatmap) and the width of the link reflect the strength of association. Using PubAtlas hence lets us identify the disease pairs in a disease class that have known former associations through literature, providing us with a basis for validating the class as well as distilling its basic characteristics.

## Acknowledgements

A.A. and M.D.D. acknowledge financial support by the Spanish government through grant FIS2015-38266. M.D.D. also acknowledges financial support from the Spanish program Juan de la Cierva (IJCI-2014-20225). A.A. also acknowledges partial financial support from the European Commission FET-Proactive project MULTIPLEX (Grant No. 317532), ICREA Academia and James S. McDonnell Foundation. A.H. and A.S. acknowledge the financial support by National Institutes of Health (NIH) grants P50-HG004233-CEGS, MapGen grant (U01HL108630) and P01 HL083069, U01 HL065899, P01 HL105339, R01HL111759, and RC HL10154301. The authors acknowledge Joseph Loscalzo for his critical reading of the manuscript and valuable suggestions, and Emanuela De Domenico for fruitful discussions.

## Author Contributions

A.H. and M.D.D. contributed equally to this work. All authors designed the study. M.D.D. and A.H. prepared the data and performed the numerical analysis. All authors wrote and approved the manuscript.

## Footnotes

1 World Wide Web URL: http://omim.org/

2 World Wide Web URL: http://www.pubatlas.org/

